# The assessment of fishery status depends on the condition of fish habitats

**DOI:** 10.1101/233478

**Authors:** Christopher J. Brown, Andrew Broadley, Fernanda Adame, Trevor A. Branch, Mischa Turschwell, Rod M. Connolly

## Abstract

At the crux of the debate over the global sustainability of fisheries is what society must do prevent overexploitation of fisheries and aid recovery of fisheries that have historically been overexploited. The focus of debates has been on controlling fishing pressure and assessments have not considered that stock production may be affected by changes in fish habitat. Fish habitats are being modified by climate change, built infrastructure, destructive fishing practices and pollution. We conceptualise how the classification of stock status can be biased by habitat change. Habitat loss can result in either overly optimistic or overly conservative assessment of stock status. The classification of stock status depends on how habitat affects fish demography and what reference points management uses to assess status. Nearly half of the 418 stocks in a global stock assessment database use seagrass, mangroves, coral reefs and macroalgae, habitats that have well documented trends. There is also considerable circumstantial evidence that habitat change has contributed to overexploitation or enhanced production of data-poor fisheries, like inland and subsistence fisheries. Globally many habitats are in decline, so the role of habitat should be considered when assessing the global status of fisheries. New methods and global databases of habitat trends, and use of habitats by fishery species are required to properly attribute the causes of decline in fisheries, and are likely to raise the profile of habitat protection as an important complementary aim for fisheries management.

## Introduction

Thirty-one per cent of marine fish stocks globally are overexploited, meaning that their potential productivity is lower than what could be supported if fishing pressure was reduced (FAO 2016). Given the large number of extinct and threatened freshwater fish species, freshwater fisheries are likely to be faring worse, although incomplete and geographically fragmented data make determinations of the current status of most inland fisheries difficult at present (Welcomme 2008, Welcomme et al. 2010). Recovering overexploited fisheries would bring significant economic and social benefits to humanity (Costello et al. 2016). Many fisheries support the livelihoods of people with few other sources of protein (Hall et al. 2013) and income from fisheries can be a major contributor to social wellbeing in coastal and inland communities (FAO 2016). In some parts of the world management regulations have successfully reduced the capacity of fishing fleets and reduced fishing pressure to levels that should enable stock recovery (Worm et al. 2009, Bell et al. 2017, Rosenberg et al. 2017). To date, the debate over the global status of fish stocks has focussed on overfishing as the primary cause of declines in fish stock production (e.g. Worm et al. 2009, Branch et al. 2011, Pauly et al. 2013). However, many well-managed fish populations are not recovering as expected (Neubauer et al. 2013, Szuwalski and Thorson 2017), suggesting other causes have contributed to declines in stock productivity. This contention is supported by evidence that many fish populations are driven by unpredictable regime changes (Vert-pre et al. 2013) and over time the productivity of many fish stocks has been declining (Britten et al. 2016).

One mechanism for unexplained changes in stock production is alterations in suitable habitat. Globally, marine and freshwater habitats are also facing considerable changes, predominantly degradation and areal loss (e.g. Welcomme 2008, Waycottetal. 2009, Ye et al. 2013, Hamilton and Casey 2016), although habitat for some species may be improving and expanding (Brown and Trebilco 2014, Claisse et al. 2014). Here we define “fish habitat” as the scenopoetic niche of aquatic species (Hutchinson 1978), which for a given species is the set of physical, chemical and biological variables that define where an organism lives. The scenopoetic niche is distinguished from the bionomic niche, which is the resources that an organism consumes (Hutchinson 1978). While there have been global-scale models of the effects of the bionomic niche on the status of fisheries (Christensen et al. 2014), global assessments of fishery status have not accounted for habitat.

There are numerous local examples that show fishery status has been affected by habitat change. For instance, hypoxia and macrophyte loss affect catches of crustacean (Breitburg et al. 2009, Loneragan et al. 2013) and bivalve fisheries (Orth et al. 2006b); nursery habitat loss limits recruitment of adults to fisheries (Beck et al. 2001, Sundblad et al. 2014); the status of many salmon fisheries has been degraded by loss of stream habitat (Gregory and Bisson 1997); and many of Europe’s major ocean fisheries depend on threatened coastal habitats (Seitz et al. 2014). Across many stocks, the types of habitats used by fish populations are an important predictor of their long-term dynamics (Britten et al. 2016, Szuwalski and Thorson 2017), implicating habitat in unexplained failure to recover. With systematic degradation of fish habitats, global assessments of stock status will overestimate the ability of fisheries management to improve the fish stock status. Surprisingly, given the long history of habitat studies, knowledge of which fisheries species are dependent on threatened habitats is incomplete even for relatively well studied habitats like mangroves (Lee 2004, Sheaves 2017) and fish habitats along Europe’s coasts (Seitz et al. 2014).

Here we argue that global assessments of fishery status are incomplete and potentially misleading without also considering habitat change. First, we review how habitat change could affect the population parameters used to determine the status of fisheries. Then we demonstrate that fisheries reference points are sensitive to habitat change using a simple model that captures the key dynamics in fisheries stock assessments. We then examine habitat use by fisheries stocks listed in the most comprehensive global database of fisheries stock assessments, the RAM Legacy database (Ricard et al. 2012). This database has been used in numerous studies to support global assessments of stock status, relative to historical fishing pressure (Worm et al. 2009, Hutchings et al. 2010, Branch et al. 2011, Costello et al. 2012, Thorson et al. 2012a, Costello et al. 2016). If there is evidence that habitat change affects a large proportion of stocks in this database, then habitat change is likely to be a globally significant driver of fisheries status. We also consider trends in other stocks with less data. Finally, because the quantitative consideration of habitat status in global stock reports remains elusive, we suggest avenues for including fish habitats in global stock assessments.

### Effects of habitat change on population demography

There are numerous mechanisms by which habitat change can affect the demography of fish populations (Vasconcelos et al. 2013). Here we describe the eco-physiological processes by which habitat change can affected population parameters under the broad categories of *carrying capacity*, *population growth rate* and *catchability* (Table 1).

**Table 1.** Examples of effects of humans on habitat change and how habitat change may affect demographic parameters and catchability.

| Parameter | Species | Habitat | Description | Reference |
| --- | --- | --- | --- | --- |
| Population growth rates | Bumphead Parrotfish (*Bolbometopon muricatum*) | Pacific coral reefs | Sedimentation causes loss of recruitment habitat | Hamilton et al. (2017) |
| Population growth rates | Coral Trout (*Plectropomus maculatus*) and Stripey Snapper (*Lutjanus carponotatus*) | Great Barrier Reef | Recruits prefer live corals of specific genera and these recruitment hotspots can enhance adult density locally | Wen et al. (2013a), Wen et al. (2013b) |
| Population growth rates | Barramundi (*Lates calcarifer*) | River, estuaries, and floodplain wetlands of Northern Australia | Restricted access to floodplain wetlands reduces food availability | Jardine et al. (2012) |
| Population growth rates | Eurasian Perch (*Perca fluviatilis*), Pikeperch (*Sander lucioperca*) | Nearshore regions with specific biophysical conditions | Availability of habitat for juveniles strongly related to regional adult abundance | Sundblad et al. (2014) |
| Population growth rates | Coho Salmon (*Oncorhynchus kisutch*) | Streams and rivers | Increasing agricultural and urban land use in catchments related to lower rates of recruitment to adult population. | Bradford and Irvine (2000) |
| Population growth rates | Atlantic Bluefin Tuna (*Thunnus thynnus*) | Pelagic zone, Gulf of Mexico | Ocean warming predicted to shift suitable habitat for spawning and larvae to earlier in spring | Muhling et al. (2011), Muhling et al. (2017) |
| Carrying capacity/population growth | Brook Trout (*Salvelinus fontinalis*) | Lakes (Ontario) | Egg survival determined by competition for spawning sites (density dependent) and habitat quality<br>(density independent) | Blanchfield and Ridgway (2005) |
| Carrying capacity | Coho Salmon<br>(*Oncorhynchus kisutch*) | Stream and lake beds | Loss of stream and lake habitat for coho eggs and juveniles has likely reduced adult recruitment, because females compete for space to spawn | Berghe and Gross (1989), Beechie et al. (1994) |
| Carrying capacity | King George Whiting<br>(*Sillaginodes punctata*) | Shallow seagrass | Loss of seagrass reduces larval recruitment and feeding opportunities | Connolly et al. (1999) |
| Catchability | Brown Shrimp<br>(*Farfantepenaeus aztecus*) | Demersal, soft sediment | Hypoxia causes aggregation of shrimp in areas with higher finfish abundance | Craig (2012) |
| Catchability | Spanner Crab<br>(*Ranina ranina*) | Benthic soft sediment | Warming temperatures stimulate seasonal migration that enhances catchability | Spencer et al. (in review) |
| Catchability | Giant Mud Crab<br>(*Scylla serrata*) | Mangrove-lined estuary | Increased river flow into estuary stimulates migration into fished areas | Loneragan (1999) |

The population’s carrying capacity is determined by the strength of intra- and inter-specific density dependence. If increasing availability of certain habitats weakens density dependence, then loss of those habitats will reduce the population’s carrying capacity. For instance, survival of salmonid eggs to juvenile stages can be strongly density dependent due to competition among adults for space to lay eggs (Berghe and Gross 1989) and competition among newly emerged juveniles (Milner et al. 2003). The available area of appropriate stream habitat will therefore put an upper limit on adult abundance. The strong dependence between habitat area and juvenile production predicts large declines in carrying capacity in many modified catchments, such as a >30% decline in the capacity of river basins in Washington State to produce Coho Salmon (*Oncorhynchus kisutch*) smolt (Beechie et al. 1994).

Habitat change may affect the population growth rate if habitat influences individual growth, survival of individuals at any life-stage, or spawning production per individual. For instance, the presence of mangroves near to coral reefs can enhance the biomass of fisheries species that live on coral reefs as adults, because juveniles use mangrove habitat (Mumby et al. 2004). Mangroves have been hypothesised to benefit juvenile fish by providing food for juveniles and by providing shelter from predation (Nagelkerken 2009). In many cases, the decision of juvenile fish to leave mangroves and migrate to reefs may be balancing a life-history trade-off between survival and growth. For instance, growth of juvenile French Grunt (*Haemulon flavolineatum*) is slower in mangroves than on reefs, but mortality rates from predation are also lower in mangroves than reefs (Grol et al. 2011). Thus, loss of mangrove habitat will affect the ability of French Grunt to balance a life-history trade-off between growth and survival, and ultimately will hinder recruitment to the adult population.

Specific oceanographic conditions are also important determinants of habitats, and changes in oceanographic conditions can affect population growth rates. For instance, spawning of the three bluefin tuna species (*Thunnus maccoyii*, *T. orientalis*, *T. thynnus*) is largely confined to tropical oligotrophic seas during seasons when ocean temperatures are between 22°C to 29°C (Muhling et al. 2017). The spawning of bluefin species in unproductive seas is an enigma, because their larvae have high energy requirements and are prone to dying from starvation (Muhling et al. 2017). Their spawning locations may be an adaptation to help their larvae avoid predators and competitors (Bakun 2013, Ciannelli et al. 2014) and ensure that their larvae experience optimal temperatures for their growth (Muhling et al. 2017). Bluefin tuna larvae grow faster in warmer waters, and individuals with higher growth rates have higher survival (Muhling et al. 2017). Warmer years have thus been shown to predict higher recruitment of juveniles for some tuna stocks, but only within their preferred temperature range, and therefore tuna recruitment success may be vulnerable to ocean warming in the future (Muhling et al. 2017).

Habitat change may affect the catchability of fish stocks by making fish more or less available to fishers. For instance, hypoxia may cause shrimp and fish to aggregate in a smaller area, thus making them easier to catch (Craig 2012). Migrations that increase the catchability of fished species can also be tied to physical habitat characteristics. Mud crab (*Scylla serrata*) catch is higher in years of greater summer rainfall, because river flow stimulates downstream movement that may increase their catchability in the lower estuary and bays (Loneragan 1999). Undetected increases in catchability may lead stock assessments to overestimate biomass and contribute to stock collapse when catch-per-unit effort is used as an index of biomass (Rose and Kulka 1999). Catchability changes caused by to habitat change may therefore bias the determination of stock status.

The relationship between habitat and demography can be more complex than habitat change affecting a single demographic parameter: there can also be interactions between demographic parameters. The quantity and quality of stream beds for Brook Trout (*Salvelinus fontinalis*) to lay their eggs may affect both the population’s carrying capacity and intrinsic population growth rate. Brook trout compete for space to spawn in gravel on stream beds, and superimposition of egg laying results in declines in the rate of survival of eggs as adult density increases (Blanchfield and Ridgway 2005). Thus, the available area of suitable stream beds affects the Brook Trout’s carrying capacity. Different stream beds also vary in their habitat quality: streams with seepage of ground water have higher egg survival (Blanchfield and Ridgway 2005). The availability of nesting sites with high groundwater seepage therefore likely affects the intrinsic population growth rate.

### Model for how habitat change can affect fishery status

Here we modify the Schaefer model to demonstrate how undetected changes in habitat could result in incorrect classification of fishery status. The Schaefer model is the logistic model of population growth with harvesting added. The status of a fishery is determined relative to the reference points of the biomass at maximum sustainable yield (*B_MSY_*,) and the fishing mortality rate at maximum sustainable yield, *F_MSY_* (Mace 2001). The maximum sustainable yield (MSY) is a theoretical value that has traditionally been the basis of management targets, but increasingly is seen as a limit reference point for precautionary fishing, to ensure that biomass should remain above *B_MSY_* (Mace 2001).

Fish stocks that grow faster, have higher survival, or are more fecund will tend to produce higher MSY values and be exploited at a higher *F_MSY_* rate. For biomass-based reference points, a stock is said to be under-fished if biomass is well above Bmsy, sustainably fished if near *B_MSY_* and overfished if biomass is well below *B_MSY_* (e.g. Ye et al. 2013). For fishing rates, if fishing mortality rate is greater than *F_MSY_*, overfishing is occurring; while if fishing mortality is lower than Fmsy, overfishing is not occurring and biomass will tend to increase to B_MSY_. Stocks are classified as recovering if they are overfished but overfishing is not occurring (F<*F_MSY_*). Note that under the logistic model, and for age-structured assessments, biomass can increase if fishing mortality is slightly more than Fmsy and biomass is below *B_MSY_*, because of density-dependent effects (Hilborn et al. 2014).

The reference points are determined by two parameters, the intrinsic rate of population increase (*r*) and the unfished biomass (*B*_0_), also known as the “carrying capacity” (*K*) (Table 1). In the logistic model we used in our illustration, *B_MSY_* is obtained at 50% of *B*_0_, but in more complex age-structured stock assessments, *B_MSY_* is on average at 40% of *B*_0_ (Thorson et al. 2012b). Habitat change could also affect the catchability (*q*) of a fish stock, which measures the relation between biomass and catch per unit effort (CPUE): biomass = qxcatch/effort While changes in catchability (*q*) do not directly affect the reference points, they can bias estimates of fishing mortality rate, because the relationship between fishing effort and fishing mortality (catch/biomass) hinges on the catchability (Walters and Maguire 1996).

We model three fisheries that are managed using reference points set on 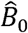, 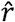 and 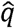, where the hat indicates the parameter is an estimate, which initially is set equal to the true values for the population. We run simulations where the true value of one of these parameters at a time changes to represent a change in habitat quality or quantity (see Table 1 for examples). We assumed an exponential decline of the true population parameters *r* and *B*_0_ over 50 years to half their initial values and a logistic increase in *q* to twice its initial value. For each type of habitat decline we modelled two management scenarios where the manager targets either 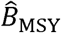 or 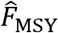 In this way, the assumed (“estimated”) values of parameters slowly diverge from the true values over time. While it is straightforward to solve for the difference between the reference points and their true values at a time, we conduct simulations here to provide a more tangible representation of how stock status would change over time when habitat is dynamic. In simulations, the manager sets fishing mortality to target a reference point, but is slow to respond to changes in stock status (e.g. Brown et al. 2012). Thus fishing mortality at a time becomes (Thorson et al. 2013):

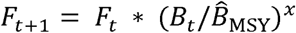

for the scenarios where 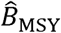 was a target and

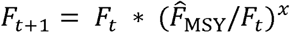

where 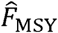 was the target. Parameter *x* controls the rate of response and we set it to 0.2 here following Thorson et al. (2013). We plot the results of simulations on “Kobe” plots, which are often used to classify the status of multiple stocks on the basis of *B_MSY_* and *F_MSY_* (e.g. Worm et al. 2009, Costello et al. 2016). Here we modify the stock status plots so that the represent status relative to the biased estimates of 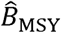 or 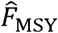 used by the manager, but we add guidelines for *B_MSY_* and *F_MSY_* as they would be estimated in the final year of simulation. Changes in catchability did not affect the estimates of the reference points, but we allowed them to cause estimation bias such that the managers assumption for the value of *F_t_* was underestimated if catchability increased. Stocks with regular assessment updates would likely result in estimated values changing over time to become closer to the true values.

We found that the effect of habitat change on stock status depends on how habitat controls population dynamics and which reference point was used in fisheries management. If habitat change affected carrying capacity, then the biomass based reference points (*B_MSY_*) were affected. Halving of the carrying capacity also required a halving of *B_MSY_*, so manager’s target of 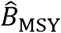 was twice what it should be. If the manager targeted 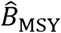 (Fig 1d) the stock was classified as below target levels due to low biomass, but, counter-intuitively, 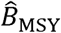 is at 2*B_MSY_*, which is the B_0_. The only way of attaining 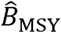 was thus to set fishing mortality rate to zero, resulting in an economic collapse of the fishery. If the manager targeted 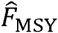 (Fig 1d) when the carrying capacity declined the stock was classified as below the biomass reference point, but on target for the mortality reference point.

**Figure 1.**
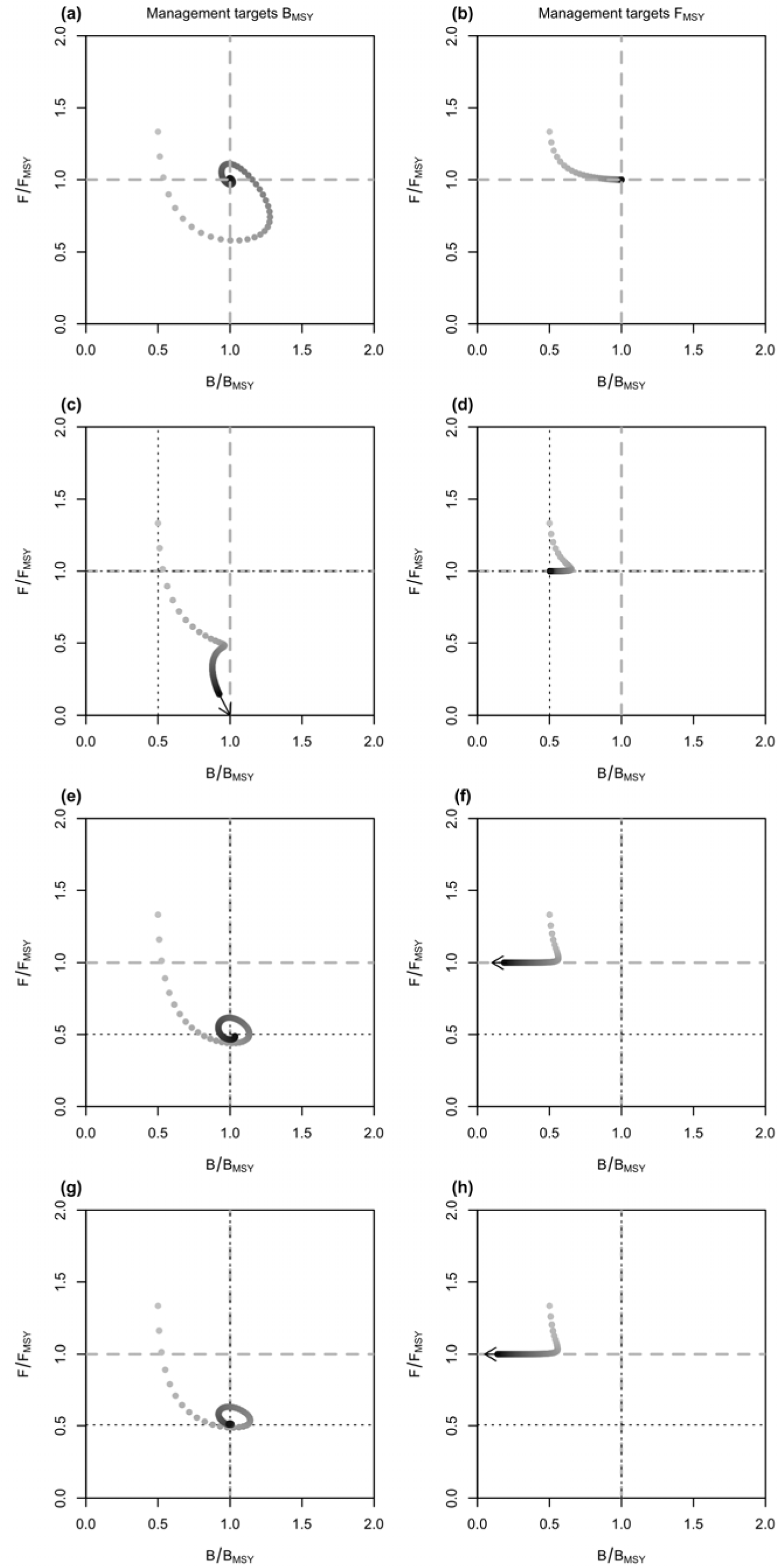
Stock status plots showing the effects of declines in habitat on the status of a theoretical fish stock. The points show biomass and fishing mortality rate for a theoretical stock over different years. starting in an overfished state fton left cmadranf) and moving toward a management target. Darker points are later years. The gray dashed lines show the management reference point if habitat change is not considered and their intersection represents the theoretical management target for a sustainably fished stock. The black dotted lines show where the reference points would shift to if habitat change was considered in a stock assessment. Arrows show trajectories for stocks that have not reached equilibrium. Simulations are given for: no change in demographic parameters (a, b), a decline in intrinsic population growth (c, d), a decline in carrying capacity (e, f) and an increase in catchability (g, h), a, c, e, g show a manager that targets 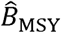 and b, d, f, h show a manager that targets 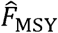.

If habitat change affected the intrinsic population growth rate then exploitation rate based reference points needed to be updated, but the biomass based reference points remained correct (Fig 1e, f). When management targeted 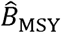 the stock was assessed as being underexploited in terms of 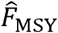 but was on target for *B_MSY_*. If management targeted 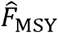 the stock was classified as underfished, resulting in its eventual extirpation (Fig 1f).

Finally, changes in catchability may cause errors in estimating exploitation rate if they are not accounted for in stock assessment models. If catchability increases, then exploitation rate may be underestimated. Thus, the effect of a catchability change on stock status (Fig 1g, h) was similar to the effect of a decline in the intrinsic population growth rate (Fig 1e, f).

To summarize, if habitat change affects a state parameter (carrying capacity) then biomass based reference points will be overly conservative and rate based reference points unaffected. If habitat change affects a rate parameter (*r* or *q*), then biomass based reference points will be unaffected but rate based reference points overly risky.

### Global trends in habitats and the status of fisheries

We reviewed habitat usage by stocks in a global stock assessment database and considered how trends in some key habitats may affect the status of those fisheries. Then we consider the effects of trends in stocks without stock assessments and for stocks that use habitat that are difficult to monitor.

#### Stocks in the RAM Legacy database

We reviewed the habitat usage of 418 marine stocks listed in the most comprehensive global stock assessment database: the RAM Legacy Stock Assessment Database (RAML; Ricard et al. 2012), available at http://ramlegacy.org/. The RAM Legacy database has been widely used to support global scale stock assessments (e.g. Worm et al. 2009, Thorson et al. 2012a) and analysis of the drivers of stock dynamics (e.g. Vert-pre et al. 2013, Szuwalski et al. 2015, Szuwalski and Thorson 2017). The RAML database has a well-known geographic bias towards temperate regions that have the scientific capacity to conduct stock assessments (Ricard et al. 2012). Numerous analyses have addressed this bias, by creating predictive models of stock status that extrapolate from the RAM Legacy database to unassessed fisheries (Costello et al. 2012, Thorson et al. 2012a). Therefore, habitat dependencies that exist in the RAM Legacy stocks are also likely to influence the status of fish stocks as estimated in many global assessments of stock status.

The review of fish habitat associations aimed to comprehensively document associations for all species in the RAM Legacy database at all life-history stages (including eggs, larvae, juvenile, adult and spawning). We searched the literature for studies that documented the habitat associations for each species listed in the RAM Legacy database. Associations were defined as any observation and include direct observation (e.g. divers), fishery catch in certain habitats, catch surveys in certain habitats and electronic tagging studies. We recorded the depth of observation, the habitat type. Habitat was classified first as whether it was physiochemical (e.g. temperature range for pelagic species) or hard substrate. Hard substrate categories included the non-exclusive categories: coral, macroalgae, mangroves, mud, reef, rock, sand, seagrass and soft-sediment. In total 7156 observations of species’ life-stages associating with particular habitats were identified in 137 references. Further details on this “fishscape” database are available at “https://github.com/cbrown5/fishscape” and in the Supporting Information.

Here we focussed on habitats where large scale regional and global trends have been documented (Table 2). These included nearshore habitats of seagrass (Waycott et al. 2009) and mangroves (Duke et al. 2007, Hamilton and Casey 2016), tropical coral reefs (De’ath et al. 2012, Perry et al. 2013, Jackson et al. 2014) and macroalgae, specifically kelp (Krumhansl et al. 2016). Seagrass and coral reefs are in decline across a large number of regions (Waycott et al. 2009, De’ath et al. 2012, Jackson et al. 2014). Kelp and mangroves show considerable regional variability, with some regions showing increases but a majority showing declines (Hamilton and Casey 2016, Krumhansl et al. 2016)(Table 2). Regardless of the variability, all these habitats exhibit significant trends that are be expected to affect the status of dependent fish stocks.

**Table 2.** Evidence of global declines in fish habitats and their potential to affect assessments of the status of fisheries. Habitat usage was reviewed for all fish species in the RAM Legacy database (Ricard etal. 2012) (SI Table 1). Catch data are for 2001, the most recent year with catch data available for all populations.

| Habitat | Evidence for global habitat change | % of fish species in RAML that use this habitat | % of catch in RAML for fish stocks that use this habitat |
| --- | --- | --- | --- |
| Seagrass | Global median rate of change - 0.9% per year, 58% of reported seagrass sites are declining (Waycott et al. 2009) | 23 | 28 |
| Macroalgae | Kelp declining in 38% of ecoregions, increasing in 27% of ecoregions (Krumhansl et al. 2016) | 23.6 | 24 |
| Mangroves | Global rate of loss of 0.16 – 0.39% per year, up to 8% in some regions (2000-2012) (Hamilton and Casey 2016). | 4.6 | <1 |
| Coral reef | Great Barrier Reef: 0.53% per year decline (1985-2012) (De'ath et al. 2012). Carribean: 50% decline in carbonate production rates (Perry et al. 2013) | 15.2 | 1.4 |

We found that 49% of species in the RAM Legacy database used habitats that included seagrass, mangroves, tropical reefs and kelp, making up 46% of the catch in 2001 (not all stock assessment time-series in the database continue to recent years). Of these major habitat types (not including the other habitats comprising 54% of catch), most species relied on macroalgae, then seagrass. Coral reefs and mangroves were less frequently used, likely reflecting the bias in the RAM Legacy database toward temperate fisheries. Most regions with good assessment coverage had >50% of species with dependence on threatened habitats (Figure 2), with the exception of the Benguela Current (South-East Atlantic). Global assessments of fisheries status are often conducted on the basis of the Food and Agricultural Organisation’s (FAO) major fisheries regions (e.g. Costello et al. 2012, Thorson et al. 2012a) and we found that >20% of stocks in most FAO regions are associated with one of the globally threatened habitat types.

**Figure 2.**
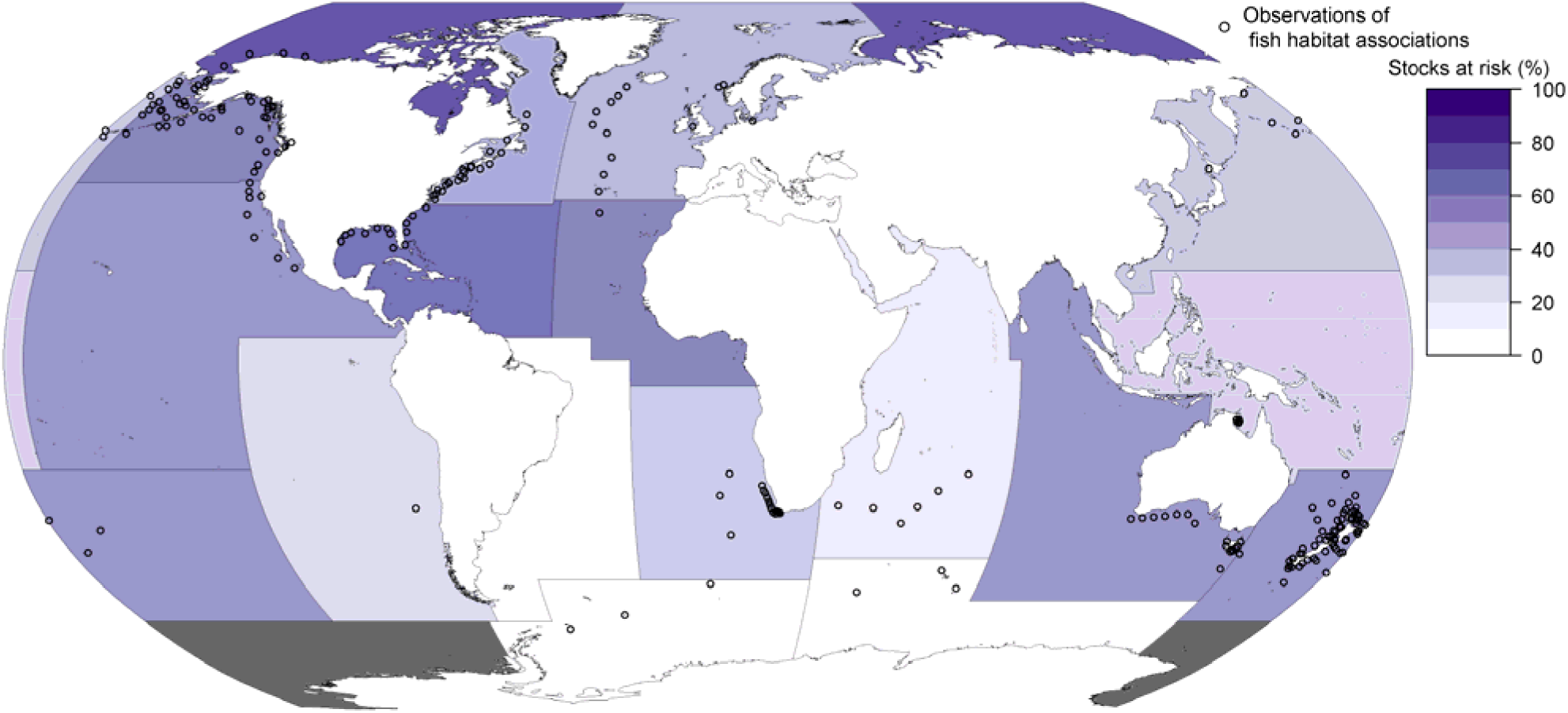
The Food and Agricultural Administration’s major fisheries areas and the per cent of stocks from a global stock assessment database that use the at-risk habitats: tropical corals, seagrass, mangroves or macroalgae.

#### Unassessed fisheries

The RAML database has some biases and importantly does not have good coverage of small-scale fisheries and tropical fisheries, further it does not include freshwater species, with the exception of diadromous fisheries. However, there is additional evidence that the global status of these data-poor fisheries is likely dependent on habitat.

First, small-scale marine fisheries, including artisanal fisheries, subsistence fisheries and recreational fisheries, are often operating in the near shore coastal zone, where the habitats showing major declines occur. Because small-scale fisheries are often limited in travel distances, they are typically close to centres of human population, which are also the epicentre of numerous threats to coastal fish habitats, like land-based stressors (Halpern et al. 2009). Threats to fish habitats in the coastal zone include land-based pollution, like increased sediment inputs from cleared land or dredging, which results in turbid waters and cause seagrass die-off (Orth et al. 2006a); nutrient inputs that can lead to algal blooms and hypoxia (Breitburg et al. 2009); clearing of mangroves for development of fish farms or coastal infrastructure (Duke et al. 2007) coastal armouring (Dethier et al. 2016) and developments that modify hydrology (Heery et al. 2017). The limited data available for small-scale fisheries means that the link between habitat change and changes in fisheries production is poorly established, however there is strong circumstantial evidence that these fisheries are affected globally by habitat change. For instance, the fisheries operating in seagrass meadows are typically small-scale, but many of the species targeted have a strong dependence on seagrass, so the global decline in the extent of meadows is expected to have impacted these fisheries (Nordlund et al. in press).

Second, tropical fisheries are not well covered in the RAM Legacy database, but many are predicted to be overexploited (Costello et al. 2012, Costello et al. 2016). The poor status of tropical fisheries has typically been attributed to their occurrence in regions with poor capacity for management (Mora et al. 2009, Melnychuk et al. 2017). However, there is strong circumstantial evidence that habitat degradation is also an important contributor to the declining status of tropical fisheries. As an example, coastal tropical fisheries are often associated with coral reefs, mangroves and seagrass, which are in decline in many tropical locations. Many fished coral reef species have close associations with their habitats (Graham and Nash 2013), support locally important subsistence fisheries (Sale and Hixon 2014) and reef habitats are threatened by human stressors including dynamite fishing, climate change (Hoegh-Guldberg et al. 2007) and land-based pollution (Brown et al. 2017, Hamilton et al. 2017). There is regional evidence that fisheries dependent on mangroves, coral reefs and seagrass are also threatened by losses of those habitats (Aburto-Oropeza etal. 2008, Uns worth and Cullen 2010, Sale and Hixon 2014).

Finally, numerous local and regional studies have documented strong associations between freshwater fisheries and their habitats, and freshwater habitats are globally threatened by pollution, habitat destruction, barrier construction and water abstraction (Dudgeon et al. 2006, Vörösmarty et al. 2010, Reis et al. 2017). Freshwater ecosystems support globally significant fisheries and their contribution to global fisheries has likely been under-estimated in official statistics (FAO 2016, Deines et al. 2017). For instance, Lake Victoria (central Africa) once supported an important fishery for indigenous predatory cichlid fish (>100 000 tons per annum (Hecky et al. 2010). That fishery has now been largely replaced by harvesting of the introduced Nile Perch (*Lates niloticus*). Multiple stressors have been implicated in the transition between these species, but an important contributing factor was run-off of fertilizer that caused eutrophication and an increase in the turbidity of the lake. Turbid conditions gave Nile Perch a competitive advantage over native predators, because they are better adapted to turbid conditions. The domination of Nile Perch in the lake did not coincide with their introduction, but occurred decades later and coincided with eutrophication (Hecky et al. 2010).

A further important impact on freshwater habitats is the construction of barriers including dams and weirs that prevent migration of freshwater fish and also alter habitat by modifying flow regimes (Dudgeon et al. 2006). For instance, dam construction in the state of Maine reduced lake habitat accessible to diadromous fish to <3% by 1900 (Hall et al. 2011) and dams on the west coast of the USA have contributed to extinctions of nearly 1/3 of Pacific salmon populations (Gustafson et al. 2007). The global trend of increasing stressors on freshwater systems would suggest that the types of ecological changes in fish assemblages that occurred in Lake Victoria may be common to many other freshwater fisheries.

## Discussion

We have argued that the global status of fisheries is dependent on trends in aquatic habitats and thus, habitat change should be considered in global assessments of stock status. Quantifying the dependence of stock status on habitat change is an important objective, because if stocks are strongly dependent on habitat change fishery, management alone will be insufficient to prevent productivity declines or recover overexploited fisheries. Habitat gains may also mask poor fishery management.

For most fished species it is unclear how habitat change will affect population demography (Vasconcelos et al. 2013). This gap needs to be filled to enable consideration of habitat change in the global assessment of fisheries. A first step could be to model habitat change as a change in carrying capacity, because this is the typical assumption used in many fisheries models, such as those used to model the spatial effects of fisheries closures on fish catch (Walters et al. 2007, Brown et al. 2015). However, we found it difficult to identify unambiguous examples of habitat change affecting a stock’s carrying capacity and noted many examples where habitat change likely affected a stock’s intrinsic population growth or catchability. For instance, it is common to measure density of a species across different habitats, but uncommon and more empirically challenging to measure whether the demographic effects of habitat are density dependent (Vasconcelos et al. 2013). Habitat change will affect carrying capacity only if survival or reproductive success is density dependent. In the absence of detailed information on the habitat-demography link, fisheries models should consider a broader range of processes by which habitat could affect population demography. For instance, evidence we found here for habitat change affecting catchability is concerning, because undetected changes in catchability have led to assessment errors that contributed to major stock collapses (Rose and Kulka 1999).

The conceptual model suggested that the impact of habitat change on a fishery’s status depends on how habitat affects the stock’s demography, whether management of fishing pressure is effective and the type of reference point used by fisheries management. Habitat loss could drive overexploitation of an effectively managed fishery, but counter-intuitively it could also cause a well-managed fishery to become more conservative and set annual catch too low, eventually resulting in economic collapse of the fishery even though the stock could in theory support some fishing. Thus, assessments of the effects of habitat change on fisheries must also consider the specific type of management targets a fishery operates on before making recommendations for altered management targets on the basis of habitat change.

It is common to consider environmental change and regime shifts when determining reference points for fisheries on a regional scale. For instance, abundance and recruitment species for many fishery species in Alaskan waters, including Pollock (*Theragra chalcogramma*) and King Crab (*Paralithodes spp*.), show pronounced regime shifts (Szuwalski and Hollowed 2016). Fisheries assessments for these species only use time-series data from the current regime to set baselines for reference points (Szuwalski and Hollowed 2016). Regime shifts are also evident at global scales, for instance, regime shifts have been detected in about 70% of abundance time-series in the RAML database (Vert-pre et al. 2013). Given the regional importance of environmental change, such drivers should be included in global stock assessments.

While a global assessment that includes habitat is an important goal, we have not conducted such an analysis here because there are several important data gaps that must be filled. There are few databases of sufficient scale that cover trends in fish habitats. Many habitats like deep sea biogenic habitats have likely been substantially degraded (Clark et al. 2015), but difficulty in observing these types of habitats means their trends have only been quantified in a few regions. Two advances in technology are creating new opportunities for global scale synthesis of hard-to-observe habitats. First, models based on meta-analysis of experimental trawling may allow global estimation of the impacts of trawling on soft-sediment habitats (Hiddink et al. 2017). Second, video monitoring and automated image analysis are enhancing capacity to observe large areas of habitats, particularly in areas that are difficult to survey with conventional methods. (e.g. Ferrari et al. 2016).

In many regions, time-series of fisheries may predate data on habitat change, so it is difficult to determine baselines for habitats. Historical studies that found evidence of habitat change often report surprising trends in fish habitats that are counter to common beliefs (Rochette et al. 2010, McCain et al. 2016, Shelton et al. 2017). For instance, on the West Coast of the USA, analysis of seagrass area as part of previously unpublished herring egg surveys unexpectedly revealed no significant long-term trends in seagrass area despite increasing coastal stressors (Shelton et al. 2017). Historical analysis of charts dating back to the 1850s reveal that the Seine estuary may once have provided nursery habitat for up to a 25% of the English Channel’s Sole (*Solea solea*) population, but habitat degradation has reduced that to a few per cent (Rochette et al. 2010). Similarly, nearshore cod-spawning areas near river outlets have been systematically lost over time, as revealed from historical analysis of cod catches (Ames 2004).

Even where there are sufficient data on habitats, it can be difficult to detect its effects on fish populations and fisheries. Often the scale of associations between fish and their habitats is not captured in habitat data and commonly measured metrics like habitat area maybe a poor indicator of how fish use habitats. Our conceptual model indicated that accurate quantification of stock status requires knowledge of how habitat interacts with population demography, but few studies have quantified how habitat affects demography (Vasconcelos et al. 2013). Such studies are important, because the demographic effects of habitat can be counter-intuitive, such as the slower growth rates of juvenile French Grunt in mangroves, even though mangroves are a preferred habitat (Grol et al. 2011).

Finally, multi-stressors may often hide the effects of habitat loss. For instance, much of Queensland’s coastal wetlands that are important fish habitats have been lost or degraded, but there is little evidence that these trends have manifest in catch of wetland dependent species by coastal fisheries (Sheaves et al. 2014). Eutrophication from nutrient run-off may have offset the effects of habitat loss by enhancing productivity and food availability for fish (Sheaves et al. 2014). Disentangling these multiple co-trending stressors may require direct studies of changes in multiple life-history processes, that includes diet composition of adult species which may benefit from enhanced production and quantifying the relationship between wetland area and survival of juveniles that use wetlands as nurseries (Haas et al. 2004).

Protection of existing habitats may often be much more effective management practice than attempting to restore lost habitats. Even coastal and shallow water habitats, like seagrass and coral reefs, which are easily accessible, can be prohibitively expensive to restore and restoration activities often have a high failure rate (Bayraktarov et al. 2016). In many cases habitat change may also represent a regime shift that is not easily reversible. For instance, once a seagrass meadow is lost, increased sediment resuspension and decreased nutrient processing act as positive feedback loops that prevent reestablishment (Maxwell et al. 2015), even with intensive restoration efforts (Katwijk et al. 2016). There are however some highly successful exceptions of habitat restoration improving the status of fisheries, particularly for freshwater fisheries (Louhi et al. 2016). For instance, dam removal can drastically increase the available habitat for salmon fisheries and be an important determinant of their recovery from historical habitat loss and overfishing (e.g. Burroughs et al. 2010).

Habitat degradation and loss may not always degrade the productivity of fisheries, so the relationship between habitat and the status of multi-species fisheries can be complex. Insufficient data to attribute the status of individual fisheries directly to habitat change means we may have missed some important cases where habitat degradation has improved the status of a fishery. Changing habitats can enhance the productivity of some species, potentially supporting new fisheries that may offset lost production from other species (Brown and Trebilco 2014). For instance, climate change and pollution have degraded many of the world’s coral reefs, but it is hypothesised that since degraded reefs can have high algal productivity they may support productive herbivore fisheries (Rogers et al. 2015, Brown et al. 2017). Artificial habitats, like oil rigs, also have the potential to enhance fisheries production (Claisse et al. 2014). Addressing the role of novel and degraded habitats in supporting productive fisheries is particularly important in regions where people have high dependence on fishing for livelihoods and subsistence, like many places with coral reefs.

Our review shows that a high proportion of stocks use habitats that are well-documented to be changing substantially on a global scale. Further, many data-poor stocks likely also have strong dependencies on habitats that are at risk of degradation. Together these results suggest that reducing fishing pressure is necessary but in many cases not sufficient to recover fish stocks and that global assessments may have overstated the potential for recovery of fish stocks on the basis of improved management of fisheries.

## Acknowledgements

CJB was supported by a Discovery Early Career Researcher Award (DE160101207) from the Australian Research Council. TAB was supported in part by the Richard C. and Lois M. Worthington Endowed Professor in Fisheries Management.

